# Foresight is required to enforce sustainability under time-delayed biodiversity loss

**DOI:** 10.1101/144444

**Authors:** A.-S. Lafuite, C. de Mazancourt, M. Loreau

## Abstract

Natural habitat loss and fragmentation generate a time-delayed loss of species and associated ecosystem services. Since social-ecological systems (SESs) depend on a range of ecosystem services, lagged ecological dynamics may affect their long-term sustainability. Here, we investigate the role of consumption changes in sustainability enforcement, under a time-delayed ecological feedback on agricultural production. We use a stylized model that couples the dynamics of biodiversity, technology, human demography and compliance to a social norm prescribing sustainable consumption. Compliance to the sustainable norm reduces both the consumption footprint and the vulnerability of SESs to transient overshoot-and-collapse population crises. We show that the timing and interaction between social, demographic and ecological feedbacks govern the transient and long-term dynamics of the system. A sufficient level of social pressure (e.g. disapproval) applied on the unsustainable consumers leads to the stable coexistence of unsustainable and sustainable or mixed equilibria, where both defectors and conformers coexist. Under bistability conditions, increasing time delays reduces the basin of attraction of the mixed equilibrium, thus resulting in abrupt regime shifts towards unsustainable pathways. Given recent evidence of large ecological relaxation rates, such results call for farsightedness and a better understanding of lag effects when studying the sustainability of coupled SESs.

## Introduction

Early research on the interaction between human populations and their environment emphasized the need for government control in order to prevent the overexploitation of natural resources [1]. However, recent empirical evidence has shown that local communities can achieve sustainable resource use through cooperative self-governance [2]. Successful communities often establish social norms, i.e. rules of shared behavior, that protect common natural resources [3] or help achieve group interests [4]. Human behavioral change can significantly affect the dynamics of social-ecological systems (SESs) [5], and is a central aspect of their adaptability, resilience and transformability [6, 7]. The evolution of social norms fosters transformability by affecting feedbacks and drivers of SESs, potentially leading to large-scale behavioral shifts [8]. Such shifts may allow escape from social-ecological traps, i.e. persistent mismatches between the responses of people and their ecological conditions, that are undesirable from a sustainability perspective [9].

The enforcement of cooperation strongly hinges on the ecological characteristics of SESs. Previous experimental and theoretical studies have emphasized the role of resource productivity and mobility [10] as well as temporal variability [11, 12] on the robustness of cooperation. Evidence from the literature on natural resource management shows that the interaction between fast and slow ecosystem processes affects the optimal management strategy [13], while inappropriate management may reinforce undesirable feedbacks and push the SES into a social-ecological trap [14]. However, the consequences of lag effects between slow and fast social-ecological processes on the robustness of cooperation remains an open question.

Lag effects can emerge from the spatial dynamics of SESs, since land conversion and fragmentation alter spatial processes and generate time-delayed loss of species [15]. Estimates suggest that 80% of the species extinctions in the Amazon are still pending [16], which may increase the number of 20th-century extinctions in bird, mammal, and amphibian forest-specific species by 120% [17]. In European landscapes, studies find that extinctions lag well behind contemporary levels of socioeconomic pressures [18], the current number of threatened species being better explained by socio-economic indicators from the early or mid-20th century [19].

The accumulation of these extinction debts generates functioning debts [20] that postpone the negative effect of biodiversity loss on ecosystem processes. Since many of the ecosystem services that play a direct or indirect role in agricultural production depend on biodiversity [21, 22, 23, 24], current species extinction rates [25, 26, 27] do not only threaten the long-term provisioning [28, 29] and stability of ecosystem processes [30, 31, 32], they also generate a time-delayed feedback loop between humans and nature [33]. In the long run, such time-delayed biodiversity feedbacks may result in large environmental crises, i.e. overshoot-and-collapse population cycles [33], similar to the famine cycles that have been observed in extinct societies [34].

Characteristics common to the majority of modern agricultural systems were found to increase the vulnerability of SESs to such crises [33]. Among these chracteristics are a high production efficiency and a low labor share per unit of agricultural good, due to the substitution of technology (e.g. machines, fertilizers and pesticides) for human labor and ecosystem services. The resulting decoupling between human population growth and ecological dynamics can reinforce unsustainable feedbacks and lead to more natural habitat loss [35]. Shifting consumption has, however, been identified as a major strategy that could allow doubling food production while greatly reducing the environmental impacts of agriculture [36, 37]. Norm-driven consumption changes towards more environmentally-friendly agricultural goods, whose production relies more on ecosystem services and labor than on technology, may thus play a key role in ensuring the long-term sustainability of SESs at large scales. However, the magnitude of biodiversity lag effects may postpone the required behavioral changes, and push (or keep) the global SES into a social-ecological trap [35].

The aim of this article is to investigate the effects of time-delayed biodiversity loss on norm-driven sustainability enforcement. To this end, we develop a dynamical system model of an endogenously growing human population divided into norm-following and norm-violating consumers, that share a common stock of land and associated biodiversity. Rising consumption demand of the human population drives production supply and natural habitat conversion through market constraints. The model thus differs from related common-pool resource systems that only consider a constant population of harvesters and a single resource [11]. The present model builds upon previous work [33], where the growth rate of the human population depends on the consumptions of industrial and agricultural goods, as well as on the strength of the demographic transition governed by technological change. The time-delayed loss of biodiversity-dependent ecosystem services then acts as a lagged feedback on agricultural productivity that can push the system into a social-ecological trap, following an overshoot-and-collapse cycle. In the following, we present the model structure and show that allowing for consumers’ behavioral change generates bistability and the potential for regime shifts. We then explore different scenarios of social pressure and biodiversity lag effects, and conclude with a discussion of our results.

## 1 Model description

### 1.1 Coupling human demography, biodiversity and social dynamics

We model a population of consumers, whose demand for agricultural and industrial goods requires the conversion of their common natural habitat. Our SES model describes the long-term interaction between four dynamical variables (Fig.1): the human population (H), technological efficiency (T), biodiversity (B) and the proportion of sustainable consumers, hereafter “conformers” (q). Conformers, by complying to a sustainable norm prescribing the consumption of environmentally-friendly agricultural goods, reduce their footprint in terms of natural habitat destruction and long-term biodiversity loss. Total habitat is gradually converted towards agricultural and industrial lands. The remaining natural habitat supports a community of species (biodiversity) that provides a range of ecosystem services to agricultural production [31]. Loss of natural habitat leads to time-delayed species extinctions, thus reducing both the common-pool biodiversity and long-term agricultural productivity [38]. Such a lagged feedback on agricultural production can result in long-term environmental crises characterized by overshoot-and-collapse population cycles (Fig.4.c). These crises transiently reduce human welfare [33], thus threatening inter-generational equity and sustainability [39]. Since the vulnerability of SESs to lag effects increases with natural habitat destruction and biodiversity loss [33], a sufficient proportion of conformers reducing their consumption footprint may help limit land conversion while preserving the long-term sustainability of the SES.

**Figure 1:**
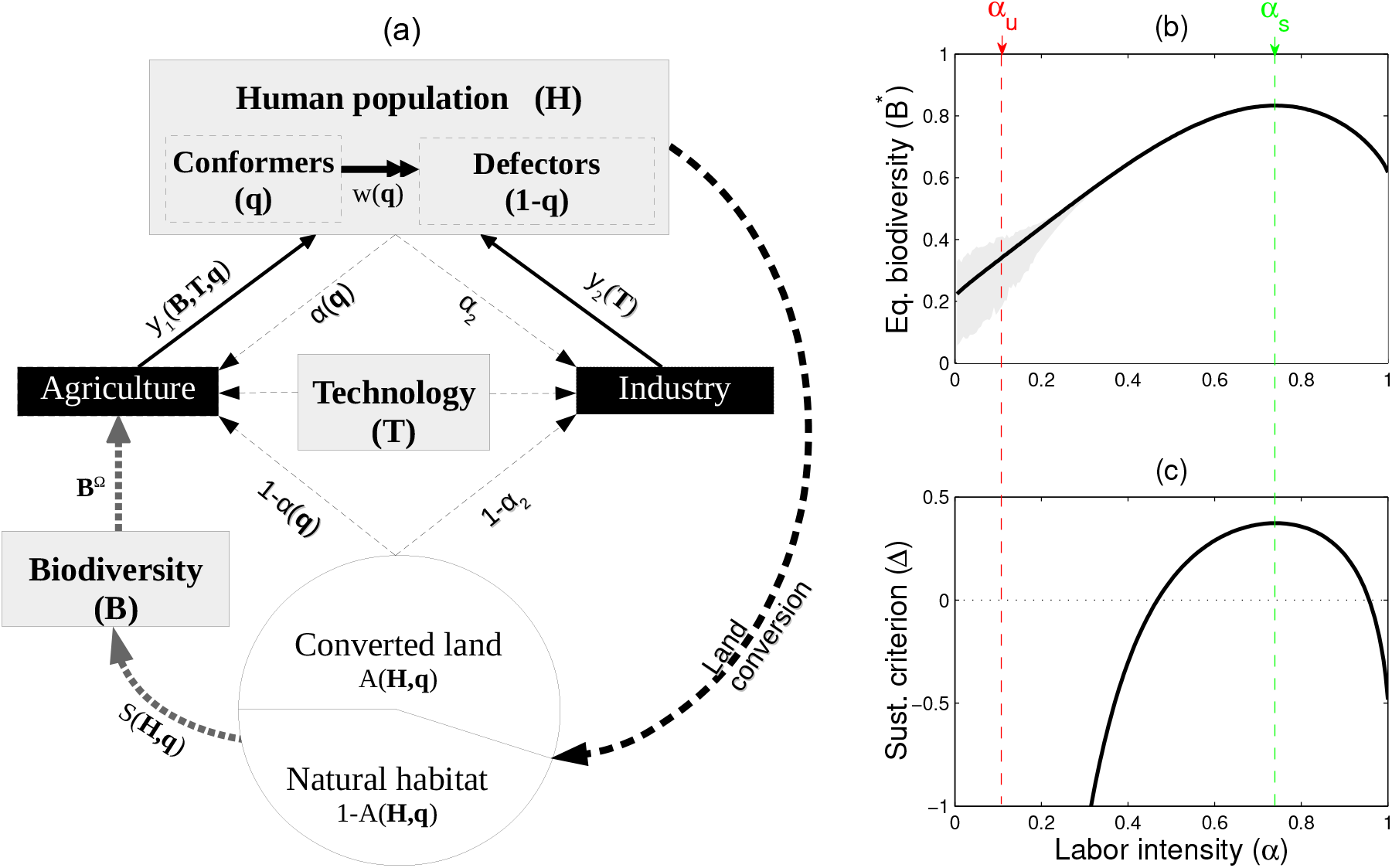
Coupling between human, social and ecological dynamics, and definition of the sustainable and unsustainable consumption norms. **(a) Model summary** Black boxes: production sectors; grey boxes: dynamical variables; dashed lines: production inputs (labor, land and technology), with *α*(*q*) being the share of labor compared to land to produce one unit of agricultural good; solid lines: *per capita* consumptions of agricultural and industrial goods, *y*_1_(B, T, q) and *y*_2_(T); grey dotted lines: ecological feedback; double arrow: social sanctioning (e.g. ostracism); circle: total land divided into converted land *A*(H, q) and natural habitat, which supports a long-term species richness *S*(H, q). All functions are explicitly defined in the main text and in Table S2 (electronic supplementary material). **(b) Effect of labor elasticity on equilibrium biodiversity**. Grey areas represent the amplitude of the transient environmental crises. **(c) Effect of labor elasticity on sustainability**. The sustainability criterion Δ is derived in [33]. Δ > 0 stands for sustainable transient trajectories, i.e. no environmental crises. The sustainability-optimal agricultural labor elasticity *α_s_* maximizes both Δ and the biodiversity at equilibrium, *B*\*. The unsustainable labor elasticity *α_u_* is chosen so that *α_u_ > α_s_* and Δ(*α_u_*) < 0.

### 1.2 A norm of sustainable consumption

The footprint of consumption goods can be related to the parameters of their production functions, and especially to the output elasticity of labor, hereafter denoted as α, and the output elasticity of land, which equals 1 − *α* under constant returns to scale. In economics, output elasticity captures the percent change in production resulting from a 1% change in an input. Under constant returns to scale, it also captures the relative share of inputs used in production. Thus, the higher *α*, the higher the labor force per unit of land used in agricultural production. Agricultural labor forces have been globally declining with the substitution of machines, fertilizers and pesticides for labor and ecosystem services, and the consequent rise in production efficiency [40] and economies of scale [41]. Conventional industrialized agricultural systems thus have lower labor elasticities *α* than environmentally-friendly systems, such as small-scale organic farming, where the substitution of labor and ecosystem services for technology is lower.

In previous work, labor elasticity has been related to the sustainability of SESs, in terms of their vulnerability to overshoot-and-collapse crises [33]. To do so, we have captured the transient dynamics of our SES by a sustainability criterion, Δ (electronic supplementary material, Table S1). This criterion depends on the difference between the ecological relaxation rate *ϵ* and the maximum growth rate of the human population, *μ*, as Δ = *ϵ* − *θμ*, where *θ* is a function of the ecological and economic parameters of the SES (electronic supplementary material, Table S1). Δ > 0 means that the ecological dynamics is fast enough compared to the human dynamics (*ϵ* > *θμ*), thus preventing transient overshoot-and-collapse crises. However, Δ < 0 means that the ecological dynamics is much slower than the human dynamics (*ϵ* < *θμ*), so that there is a high probability of experiencing transient crises. Using this criterion, we show in Lafuite & Loreau [33] that a low labor elasticity, i.e. a high substitution of natural and human capital for technology, increases the vulnerability of SESs to lag effects, while there exists an intermediate sustainability-optimal labor elasticity that maximizes both long-term biodiversity (Fig.1.a) and sustainability (Fig.1.b).

Let us define as *α_s_* the sustainability-optimal labor elasticity, and *α_u_ < α_s_* an unsustainable labor elasticity chosen such that Δ(*α_u_*) < 0 (Fig.1.a). The average agricultural labor elasticity then varies with the proportion of conformers as *α*(q) = q*α_s_* + (1 − q)*α_u_*. Through means of eco-labeling, consumers can either buy sustainable agricultural products (*α_s_*), or follow their unsustainable consumption habits and buy unsustainable agricultural products (*α_u_*). Since economic dynamics are much faster than ecological and demographic dynamics, we assume that agricultural and industrial production instantaneously follows consumers’ demand. Such a shift in agricultural production may not be met instantaneously due to inertia and production barriers [42], and farmers’ adaptability may have to be supported through adequate policy changes [43]. However, given the large time scales considered here, it seems reasonable to neglect such time delays with respect to the extent of extinction debts. Thus, in our system, a consumption shift towards sustainable goods, which, in turns, drives a shift towards more environmentally-friendly agricultural practices, may prevent environmental crises.

### 1.3 Human demography and technological change

The growth rate of human populations can be related to consumption levels [44] by capturing basic linkages between technology and human demography [45, 46, 47]. Using the same auxiliary economic model as in Lafuite & Loreau [33], we derive the *per capita* industrial and agricultural consumptions as functions of biodiversity and technological efficiency. Industrial consumption, *y*_2_ = *γ*_2_T/*T_m_*, varies with technological efficiency only, while agricultural consumption *y*_1*i*_ = *γ*_1*i*_B^Ω^T/*T_m_* of conformers (*i* = *s*) and defectors (*i* = *u*) also depends on biodiversity-dependent ecosystem services. The relationship between biodiversity and ecosystem services [31] is captured by a concave-down function of biodiversity, B^Ω^, with Ω < 1 [48]. *γ*_1*i*_ and *γ*_2_ are functions of other socio-economic parameters of the SES (electronic supplementary material, Table S1), and especially of labor elasticity, *α_i_* (*i* = {*u, s*}) and *α*_2_, respectively.

Both industrial and agricultural consumptions increase with technological efficiency, T. Technological change is assumed to follow a logistic growth at a rate *σ* towards a maximum efficiency, *T_m_*, in order to reproduce past agricultural productivity rise and current stagnation [49].

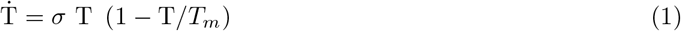

By increasing production efficiency, technological change helps counterbalancing the feedback of biodiversity loss on agricultural productivity in the short term, thus ensuring that the consumption utility of consumers does not decrease with time [33].

Following previous studies [44, 47], we then assume that the human growth rate endogenously varies with the mean agricultural and industrial consumptions, so as to increase with agricultural consumption, and decrease with industrial consumption, capturing the effect of the demographic transition.

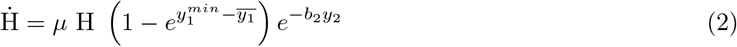

*μ* is the maximum growth rate, 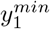 is the minimum consumption threshold, *ȳ*_1_ = q · *y*_1*s*_ + (1 − q) · *y*_1*u*_ is the average agricultural consumption, and *b*_2_ is the demographic sensitivity to industrial consumption. The strength of the demographic transition thus gradually increases with technological change and limits human population growth.

Dependence of the human growth rate on consumption levels also allows coupling human demography with social changes regarding consumption choices. Indeed, conformers do not only have a lower consumption footprint than defectors, they also have a lower agricultural consumption level. It can be shown that *y*_1*s*_ < *y*_1*u*_, meaning that for a given revenue, sustainable consumption is more costly than unsustainable consumption. As a result, conformers also have a lower reproduction rate compared to defectors. This can be interpreted as a quantity-quality trade-off in both consumption choices and the number of children, a mechanism which has been shown to partly explain the fertility reductions observed during the demographic transition [50]. Under our assumptions, shifting behaviors towards sustainable consumption habits thus reduces the growth rate of the human population, therefore increasing the sustainability of the SES.

### 1.4 Land conversion and biodiversity dynamics

Using the same auxiliary economic model as in Lafuite & Loreau [33], we derive the rate of land conversion as function of the dynamical variables of our system, under sustainable and unsustainable labor elasticities, *α_u_* and *α_s_*. For a given proportion of conformers q and human population H, converted area writes 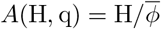, where 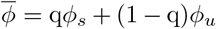 is the mean population density on converted land, and *ϕ_u_* and *ϕ_s_* are explicitly defined as functions of the economic parameters of the SES in Table S1 (electronic supplementary material). Natural habitat conversion results in time-delayed changes in species richness [51], so that the long-term species richness may be reached only after decades [16]. These extinction debts [52] are a result of many mechanisms [53] which lower the relaxation rates of communities [54]. We use a power-law species-area relationship to capture the dependence of long-term species richness on the remaining area of natural habitat [55, 56, 57, 58]. Since *A*(H, q) ∈ [0; 1], we allow the long-term species richness to vary between 1 (no habitat conversion) and 0 (all habitat is converted) by writing *S*(H, q) = (1 − *A*(H, q))^*z*^, where the slope *z* ∈ [0; 1] ensures that the function is concave-down [48]. Following experimental and theoretical results [59, 16, 60, 54], we then assume that the rate of community relaxation is proportional to the difference between current biodiversity B and long-term species richness *S*(H, q).

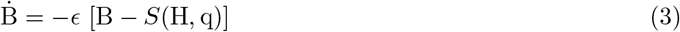

The inverse of the relaxation coefficient *ϵ* measures the time it takes to lose approximately 63% of the species that are doomed to extinction [54].

### 1.5 Social dynamics

Let us assume that the human population has identified the sustainability-optimal agricultural labor elasticity, *α_s_* (Fig.1.b and c). Restricting one’s consumption to sustainable agricultural goods has become a social norm, i.e. a shared rule of behavior. Recent studies demonstrate the importance of social norms on eating behaviors [61] and their role in shifting preferences towards healthy food [62, 63, 64]. The importance of dietary social norms is especially important in young adults [65], whose eating patterns typically become life-long habits [66]. Perception of others’ pro-environmental behavior was identified as the first step towards environmentally-friendly behavioral change [67].

Deviance from a social norm can lead to direct or indirect sanctioning from other members of the SES, be they important others or strangers [68]. Ostracism can result in social exclusion or poor reputation [69], thus decreasing the well-being of individuals. As a consequence, social pressure can reduce the well-being of defectors to the point where it becomes more profitable for them to shift behavior in order to conform to the sustainable norm. We approximate the well-being of consumers by their consumption utility, which is a function of their *per capita* agricultural and industrial consumptions. Let us denote the utility of a consumer of type *i* (*i* = {*u, s*}) as 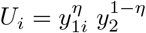, where *η* is the preference for agricultural goods. Under our assumption that *α_s_* > *α_u_*, it can be shown that the consumption utility of defectors in the absence of social pressure, *U_u_*, is higher than the consumption utility of conformers, *U_s_*. Therefore, in the absence of social pressure, defectors have no incentive to shift their habits.

Following previous studies [11, 12], we assume that social pressure decreases the utility of defectors, *U_d_* = *U_u_* − *w*(q) · *δ_U_*, so that it may become more profitable for defectors to shift their consumption and comply to the sustainable norm. The severity of the ostracism function, *w*(*q*) = *w_max_e^tηe^rηq^^*, increases with the proportion of conformers in the population, *q*, and depends on the maximum sanctioning *w_max_*, the sanctioning effectiveness threshold *t*, and the growth rate of the function, *r*. In addition to depending on the number of conformers in the community, graduated sanctioning and equity considerations leads conformers to act more strongly against defectors which consumption is the most unsustainable [3]. Thus, the lower *α_u_* and the larger the difference in consumption utilities between conformers and defectors, *δ_U_* = (*U_u_* − *U_s_*)/*U_u_*, the stronger the social pressure.

The proportion of conformers then follows a replicator dynamics [11, 12], i.e. varies both with the proportion of conformers q, and the difference between the sustainable consumption utility, *U_s_*, and the average consumption utility, *Ū* = q · *U_s_* + (1 − q) · *U_d_*.

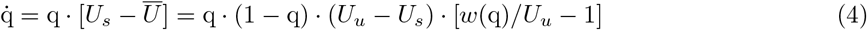

Since *U_u_* > *U_s_* in our model, a global dietary shift towards sustainable consumption (q > 0) is only possible if the severity of the social pressure is higher than the utility of defectors in the absence of social pressure, i.e. *w*(q) > *U_u_* (eq.(4)).

In the following, our focus is on the potential of consumers’ behavioral change in preventing unsustainable trajectories, i.e. overshoot-and-collapse population crises leading to biodiversity-poor equilibria in the long run [33]. We first analyze the dynamical system of equations (1), (2), (3) and (4), with a negligible ecological relaxation rate (*ϵ* = 0.1). The consequences of lag effects are explored in section 2.4.

## 2 Results

### 2.1 Social-ecological equilibria

Our SES can have two types of equilibria (*H*\*, *B*\*, *T_m_, q*\*), hereafter denoted as viable (*H*\* > 0 and *B*\* < 1) or unviable (*H*\* = 0 and *B*\* = 1), when the economic parameters do not allow the human population to maintain itself in the environment [33]. Let us denote the viable equilibria as 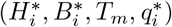, with *i* = {*u, s, c*}. Among the viable equilibria, one is unsustainable (*i = u*), i.e. only defectors persist 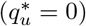 and the transient dynamics includes overshoot-and-collapse population cycles under large extinction debts (Fig. 4.c). The other two types of viable equilibria are either fully sustainable (*i* = *s*) when only conformers persist 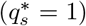, or partially sustainable (*i* = *c*) when both conformers and defectors coexist 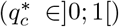. The coexistence equilibrium satisfies 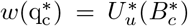, for which there is no analytical solution.

A general analytical solution for the unsustainable and fully sustainable equilibria is given in eq.(5), where the population density *ϕ_i_* and *γ*_1*i*_ (*i* = {*u, s*}) are explicitly defined as functions of the parameters of the SES in Table S1 (electronic supplementary material).

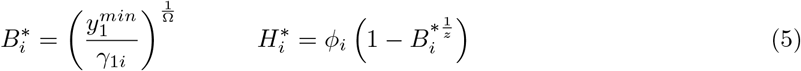

Under our assumption that *α_s_* > *α_u_*, it can be shown that *γ*_1*s*_ < *γ*_1*u*_, so that biodiversity at the sustainable equilibrium is higher than that at the unsustainable equilibrium, i.e 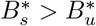. However, population density is also higher at the sustainable equilibrium, i.e. *ϕ_s_* > *ϕ_u_*, so that the human population size at equilibrium does not necessarily decrease with the proportion of conformers. Compliance to the sustainable consumption norm thus helps preserving biodiversity while not necessarily reducing the size of the human population.

### 2.2 Alternative stable states

A stability analysis of our SES model shows that two of the viable equilibria can be both stable at the same time, depending on the severity of the ostracism function compared to the consumption utility at equilibrium (electronic supplementary material, section 3). The *per capita* consumption utilities at the sustainable, unsustainable and coexistence equilibria are equal to 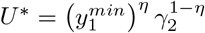. Compliance to the sustainable consumption norm thus does not reduce the long-term consumption utility.

The sustainable equilibrium is stable if the maximum ostracism *w*(1) is higher than the consumption utility that the defectors would have at the sustainable equilibrium, i.e. 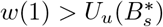, where 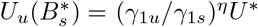. Conversely, the unsustainable equilibrium is stable if the minimum ostracism *w*(0) is lower than the consumption utility at the unsustainable equilibrium, i.e. 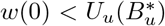 where 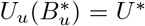 (Fig.2.a). Therefore, for intermediate consumption utilities, 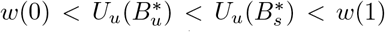, both the unsustainable and sustainable equilibria are stable ((U/S) region in Fig.2.b). For high consumption utilities, *w*(0) < *U*\* and 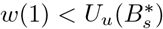, ostracism is too low to enforce norm-driven behavioral change, and the unsustainable equilibrium is the only stable equilibrium that the SES can reach ((U) region in Fig.2.b), or both the unsustainable and mixed equilibria are stable ((U/M) region in Fig.2.b). Since there is no analytical expression for the mixed equilibrium, we are not able to derive any stability condition for this bistability region. However, Fig.2.b shows that the shift between the two bistable regions (U/M) and (U/S) depends on the footprint *α_u_* of unsustainable consumption. The larger the footprint of defectors compared to conformers (*α_u_* ≪ *α_s_*), the larger the bistability region (U/M) between the mixed and unsustainable equilibria and the smaller the bistability region (U/S). Thus, the larger the required behavioral change to shift from unsustainable habits (*α_u_*) towards sustainable habits (*α_s_*), the more difficult it is to reach sustainability.

**Figure 2:**
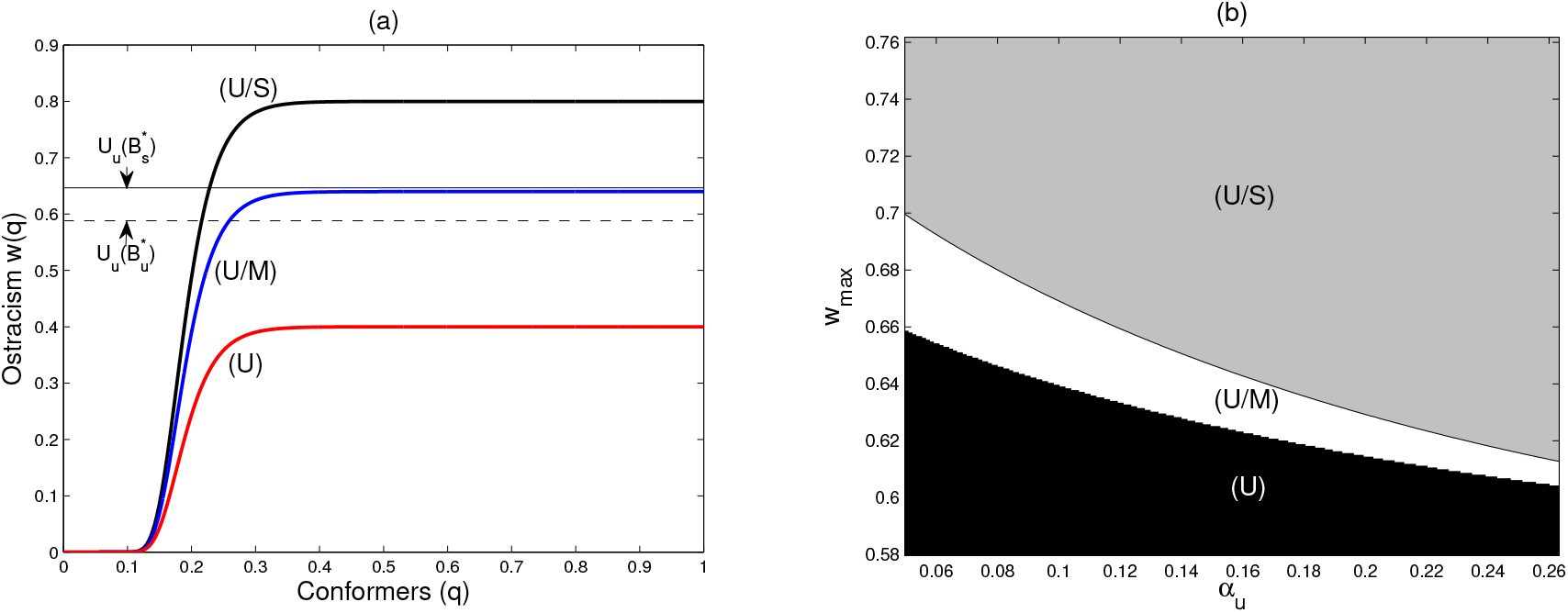
Combined effect of the ostracism parameters and the difference between sustainable and unsustainable norms on the stability of the equilibria. (a) Shape of the ostracism function for varying maximum ostracism *w_max_*. (U) Weak sanctioning (*w_max_* = 0.4) and stability of the unsustainable equilibrium only, such that 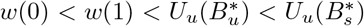; (U/M) intermediate sanctioning (*w_max_* = 0.64) and bistability of the unsustainable and the mixed equilibria; (U/S) strong sanctioning (*w_max_* = 0.8) and bistability of the unsustainable and the sustainable equilibria, such that 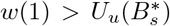 and 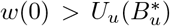. See table S1 in the electronic supplementary material for other parameter values. **(b) Stable equilibria with varying maximum ostracism** *w_max_* **and unsustainable labor elasticity** *α_u_*. Regions (U), (U/M) and (U/S) correspond to the red, blue and black curves in (a), respectively.

### 2.3 Impact of the initial state of the SES

Depending on the parameters of the SES, the size of the human population at the sustainable equilibrium can be either higher (e.g. for *T_m_* = 2) or lower (e.g. for *T_m_* = 1.8) than at the unsustainable equilibrium. Let us now consider a situation where the human population size at the sustainable equilibrium is lower than that at the unsustainable equilibrium.

Fig.3 shows that, when there is bistability ((U/M) and (U/S) panels), the sustainable and mixed equilibria are only reached in the long run when the initial proportion of conformers is high enough. The stronger the ostracism, the lower the minimum proportion of conformers required for sustainability enforcement, i.e. the larger the sustainable basin of attraction. Gradually changing social parameters may thus push an initially unsustainable SES ((U) panel in Fig.3) towards a sustainable path ((U/S) panel in Fig.3), provided that the initial social capital is large enough.

Under conditions of bistability, the type of equilibrium that will be reached in the long run thus depends on the rate of social change. In the following, we show that the rate of social change also depends on human perception of environmental changes and, in our case, biodiversity lag effects.

**Figure 3:**
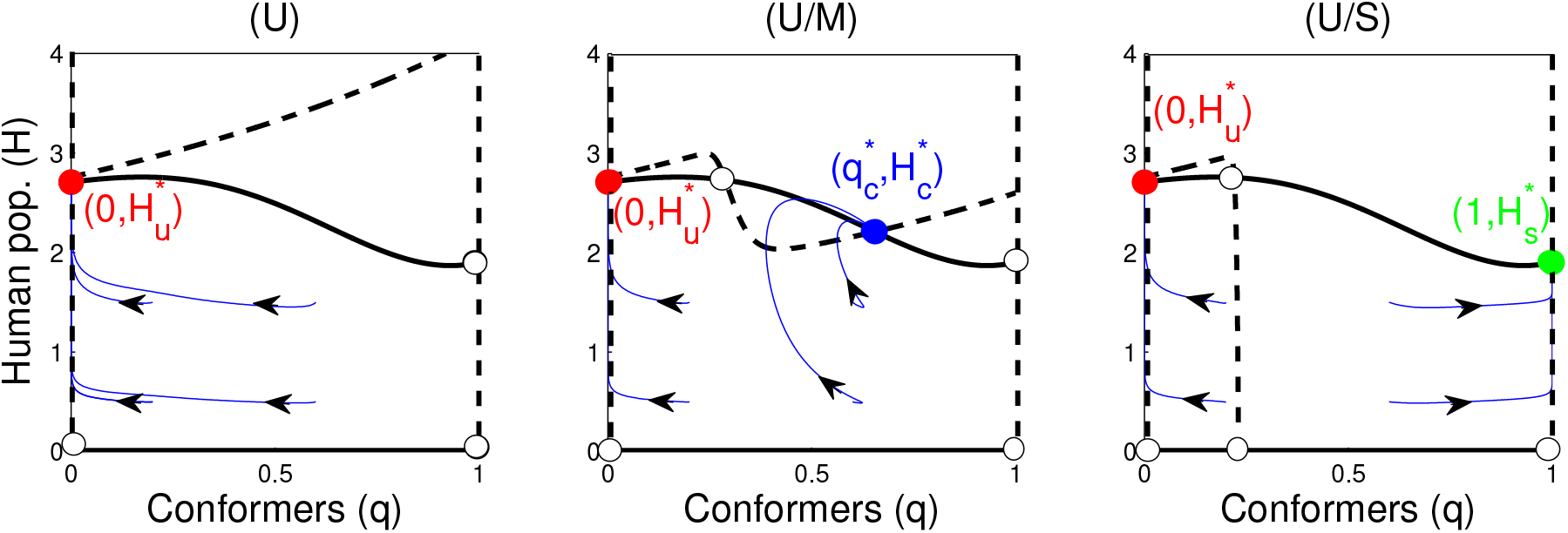
Effect of the initial conditions on the long-term equilibria, for various ostracism strengths *w_max_*. Cases (U), (U/M) and (U/S) correspond to the ostracism functions defined in Fig. 2.a. Black curve: isocline Ḣ = 0; dashed curve: isocline 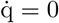; white dots: unstable equilibria; green dot: (stable) sustainable equilibrium, 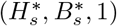; red dot: (stable) unsustainable equilibrium, 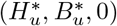; blue dot: (stable) coexistence equilibrium, 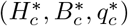; transient trajectories are represented by the blue curves. See table S1 in the electronic supplementary material for parameter values.

### 2.4 Impact of lag effects on the effectiveness of ostracism

We now explore the transient behavior of the SES with varying ecological relaxation rates, *ϵ*, for two of the initial conditions used in Fig.3, corresponding to two initial proportions of conformers q(0) = 0.2 and q(0) = 0.6, with the same human population size H(0) = 0.5. In order to better visualize transient environmental crises, we plot the null-clines and transient trajectories in the human-biodiversity phase plane (Fig.4.a). Ecological relaxation rates slow down the social dynamics by postponing the utility reduction of defectors, *U_u_*, and therefore, their consumption shift towards sustainable habits (eq.(4)). When the extinction debt is moderate, transient dynamics towards the unsustainable and mixed equilibria show environmental crises, the amplitude of which is lower for the mixed equilibrium (Fig.4.b). The sustainable trajectories do not experience any overshoot-and-collapse behavior, even for high extinction debts ((U/S) panel in Fig.4.c), which confirms the relevance of our sustainability criterion. A high extinction debt leads to very large environmental crises over the unsustainable trajectories (Fig.4.c). Moreover, in the case of bistability between the unsustainable and mixed equilibria, all trajectories now reach the unsustainable equilibrium ((U/M) panel in Fig.4.c). Large ecological relaxation rates thus result in the loss of stability of the mixed equilibrium in favor of the unsustainable equilibrium. This result suggests a shift in the dominant social-ecological feedback for increasing relaxation rates. At low relaxation rates, the ecological dynamics is fast enough for the negative effect of environmental degradation on human well-being to result in a fast enough social changes, thus reinforcing sustainable feedbacks through an efficient social ostracism. However, large extinction debts slow down the ecological dynamics and postpone the negative ecological feedback on human well-being. This reduces the efficiency of social ostracism and results in a shift of the dominant feedback towards unsustainable feedbacks, i.e. increasing consumptions and decreasing labor intensities.

**Figure 4:**
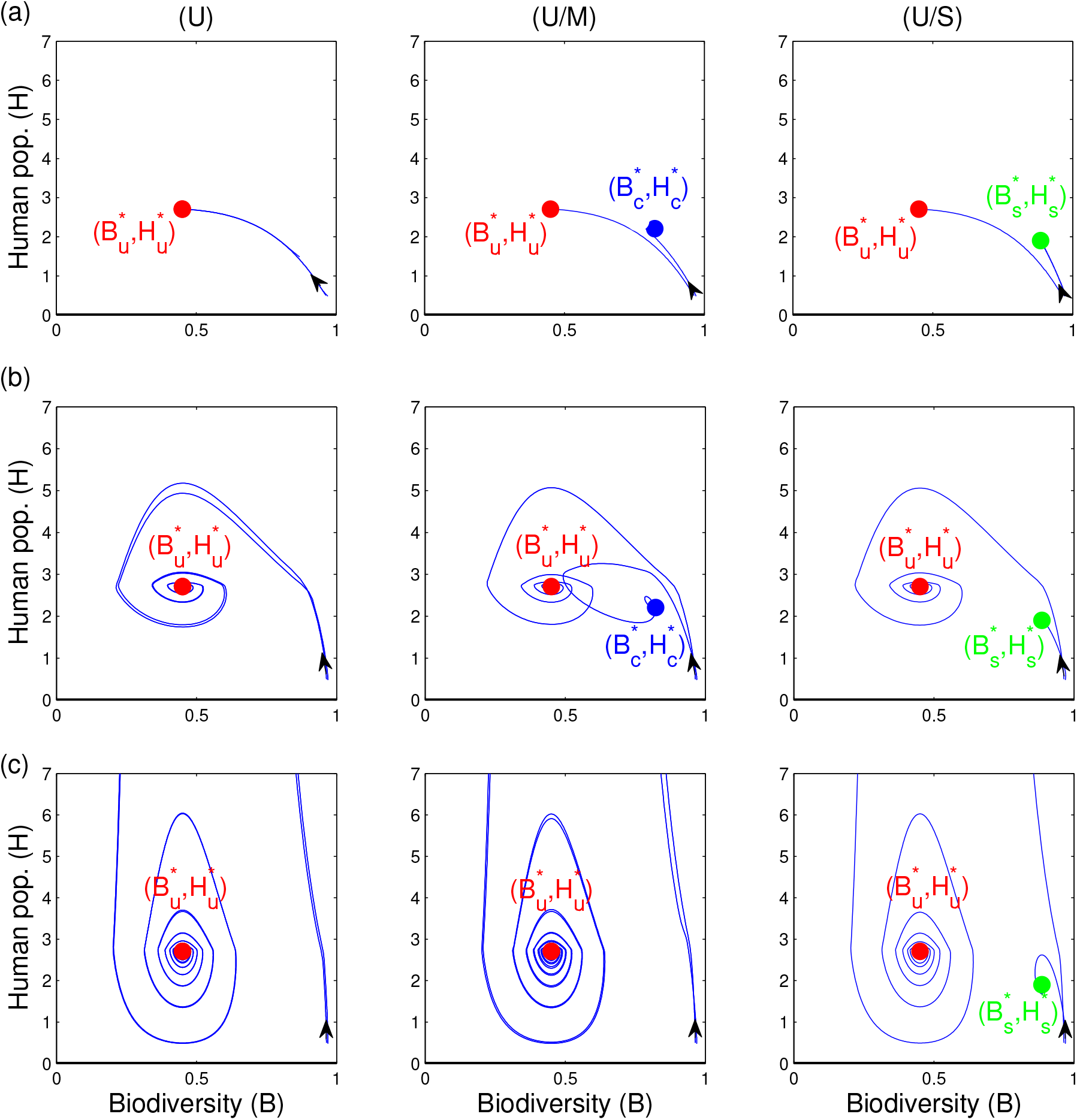
Effect of varying extinction debts e and ostracism strengths *w_max_* on transient dynamics and stability, for two initial proportions of conformers *q*(0). Cases (U), (U/M) and (U/S) correspond to the ostracism functions defined in Fig. 2.a, with similar initial conditions, i.e. H(0) = 0.5, B(0) = (1 − H(0)/*ϕ*(q))^*z*^ and q(0) = 0.2 or q(0) = 0.6. **(a) low extinction debt** (*ϵ* = 0.1); **(b) intermediate extinction debt** (*ϵ* = 0.0025); **(c) large extinction debt** (*ϵ* = 0.0005); green dot: (stable) sustainable equilibrium; red dot: (stable) unsustainable equilibrium; blue dot: (stable) mixed equilibrium; transient trajectories are represented by the blue curves. See table S1 in the electronic supplementary material for other parameter values.

Fig.5 shows the combined impact of lag effects and social ostracism on the basins of attraction of the sustainable, mixed and unsustainable equilibria. Increasing both the initial proportion of conformers and the strength of the ostracism can push an initially unsustainable SES into the basin of attraction of the sustainable or mixed equilibria (Fig.5.a). However, decreasing the ecological relaxation rate *ϵ* reduces the basin of attraction of the mixed equilibrium in favor of the unsustainable equilibrium (Fig.5.b). The stability of the mixed equilibrium appears to be much more sensitive to ecological time-lags than that of the sustainable equilibrium. Thus, moderate behavioral changes leading to a mixed equilibrium may not be robust enough to ecological lag effects. These results suggest that only important behavioral changes allowing to reach the fully sustainable equilibrium may be able to counteract the destabilizing effect of ecological time lags. The extinction debt, by postponing the consequences of environmental degradation on human well-being, thus reduces the robustness of social change and norm-driven sustainability enforcement.

**Figure 5:**
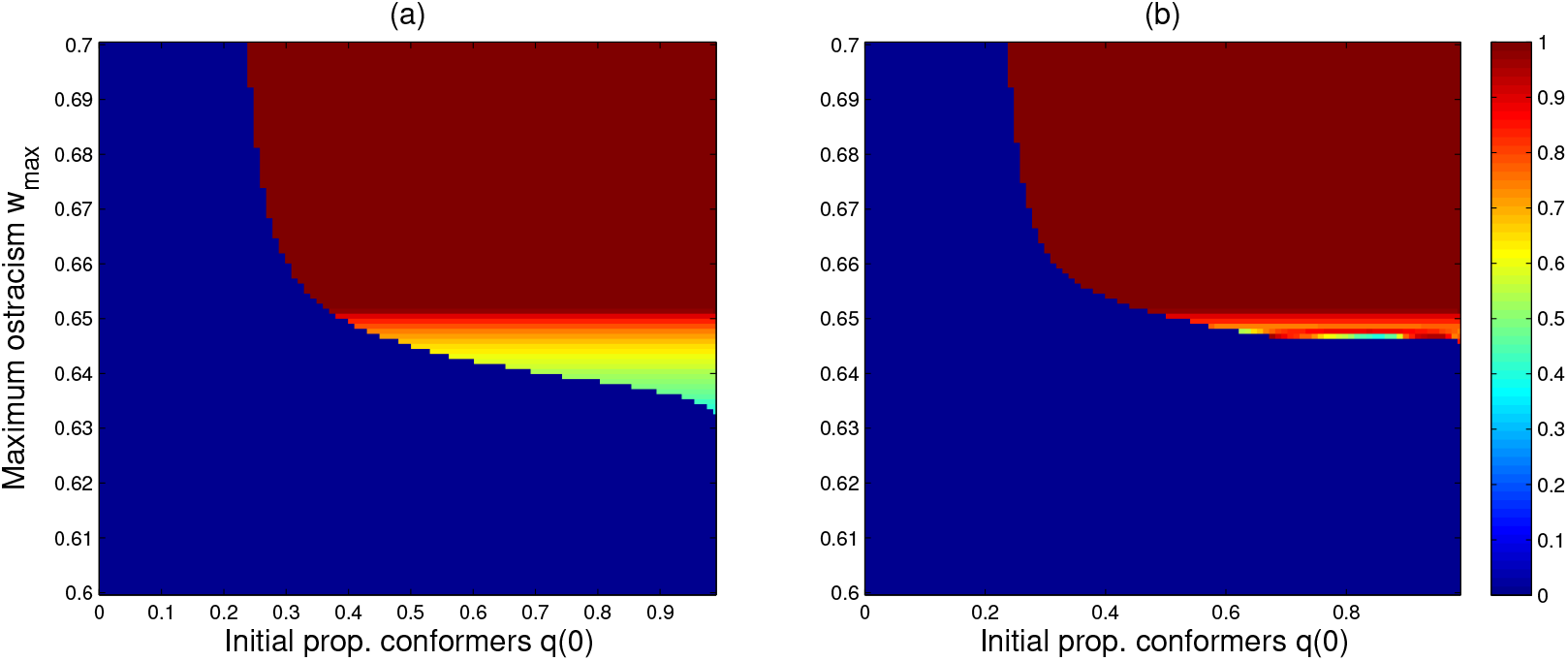
Proportion of conformers at equilibrium (*q*\*) under the combined effects of the initial proportion of conformers *q*(0) and maximum ostracism *w_max_*, for an increasing extinction debt. Other initial conditions: *H*(0) = 0.5, and *B*(0) = (1 − *H*(0)/*ϕ*(*q*(0)))^*z*^. See table S1 in the electronic supplementary material for other parameter values. **(a) low extinction debt** (*ϵ* = 0.2); **(b) high extinction debt** (*ϵ* = 0.0005). Blue color represents a population of defectors (*q*\* = 0), red color represents a population of conformers (*q*\* = 1) and yellow represents a coexistence of conformers and defectors (*q*\* ∈ [0, 1]).

## 3 Discussion and conclusions

We investigate the robustness of norm-driven sustainability enforcement, as measured by a shift towards low-footprint consumption habits. Specifically, we focus on the robustness of SESs to time-delayed biodiversity losses caused by human-driven natural habitat destruction. Lag effects in ecosystem processes are known to reinforce negative management feedbacks and potentially push SESs into social-ecological traps [13]. However, little research so far has investigated the long-term impacts of biodiversity lag effects on the sustainability of coupled SESs. Ecological studies of the anthropogenic impacts on resources or ecosystems often neglect changes in the size and behavior of the human population. Additionally, natural resources are often managed as decoupled from the ecosystems they are part of, and most socio-economic studies overlook the finiteness and physical limits of natural systems. Modeling sustainability requires accounting for the bidirectional coupling between human and natural systems [70], and especially the feedback loop between human population growth and environmental degradation [33].

In our model, this feedback loop is mediated through biodiversity-dependent ecosystem services to agricultural production. A human population exploits a shared land resource divided into natural habitat and converted agricultural and industrial lands. Natural habitat supports a community of species and provides a range of biodiversity-dependent regulatory services to agricultural production, which can itself be seen as a provisioning service. A norm of sustainable consumption is maintained through social sanctioning of unsustainable consumers. Increasing demand for sustainable consumption translates into more sustainable agricultural practices, which involve the use of a larger proportion of labor compared to land. Finally, human population growth is driven by the interaction between available agricultural resources, technological and social changes, thus adding to the growing literature modeling the interaction between human populations and their environments [34, 71].

The sustainable consumption norm is identified following Lafuite and Loreau’s (2017) sustainability criterion, which characterizes the vulnerability of an SES to transient “overshoot-and-collapse” population crises. This criterion captures the difference between the rates of ecological relaxation and human population growth, so that sustainable SESs have high enough ecological relaxation rates compared to the growth rate of their human populations. We verify here the validity of this sustainability criterion, showing that a shift towards more environmentally-friendly agricultural practices, i.e. characterized by a lower substitution of ecosystem services and labor for technology, decreases the vulnerability of SESs to transient crises. Such a global shift towards sustainable agricultural practices would require reversing current trends of land-intensive and highly mechanized agricultural production towards more labor-intensive productions, e.g. small-scale agro-ecological farms. Growing evidence suggests that diverse small-scale agro-ecological farms increase carbon sequestration, support biodiversity, rebuild soil fertility and sustain yields over time, thus securing farm livelihoods, while competing with industrial agriculture in terms of total outputs, especially under environmental stress [72].

Under a negligible ecological time delay between natural habitat loss and biodiversity erosion, full sustainability is ensured when both social sanctioning and the proportion of conformers are large enough, and when the required behavioral change to shift from unsustainable to sustainable habits is not too large. Otherwise, a minority of defectors coexists with a majority of conformers at the mixed equilibrium. When social sanctioning and/or the proportion of conformers is too low, only defectors persist at equilibrium. This unsustainable equilibrium is always stable, so that there is bistability between the unsustainable and sustainable or mixed equilibria, when these are stable. These findings echo those of Tavoni et al. [11], who used a similar non-costly social sanctioning to study cooperation enforcement in the management of a single natural resource under variable environmental conditions. However, time delays have an opposite effect to resource variability, since temporal variability tends to decrease the mean resource level, thus increasing the probability of a behavioral shift towards norm compliance. Our model differs from Tavoni et al. [11] in many aspects; first, here we focus on the interaction between various ecosystem services, especially provisioning and regulatory services, instead of a single natural resource; second, these services feed back on the dynamics of the human population that uses these services, so that the human population varies endogenously with the state of the environment; lastly, social sanctioning affects consumers’ behavior, instead of producers’. The latter feature allows us to focus on the potential of consumers’ behavioral changes in enforcing sustainability in coupled SESs.

Our study provides insights into the consequences of lag effects for norm-driven sustainability enforcement. Biodiversity loss acts as a negative feedback on human well-being, through the loss of biodiversity-dependent regulatory services to agricultural production. A time-delayed biodiversity feedback thus maintains a high utility of defectors for a longer period of time. This time lag decreases the efficiency of social ostracism, thus delaying behavioral shift. Postponing the behavioral shift of defectors towards sustainable consumption for too long can make the mixed equilibrium totally unreachable, meaning that a tipping point has been crossed in terms of human population size and habitat destruction. The lower stability of the mixed equilibrium and its propensity to regime shifts was already observed by Lade et al. [73]. Thus, under large time delays, the only way to reach sustainability is to reach the full-sustainability equilibrium, which requires much larger behavioral changes. However, given the widely observed coexistence of both conformers and defectors in small groups [74], such behavioral changes seem rather unlikely.

Moreover, theory suggests that relaxation rates are not constant, but increase with the extent of habitat destruction and fragmentation [51], thus further delaying the feedback of biodiversity-dependent ecosystem services on human societies [20]. In situations where habitat destruction leads to a strong increase in ecological relaxation rates, we would expect a decrease in sustainability, or a shift towards unsustainable development paths. An interesting extension to our work would thus be to use a spatially-explicit ecological model, in order to gain more realism regarding the temporal dynamics of ecological relaxation rates under habitat destruction, and study social-ecological regime shifts from a spatial perspective.

The emergence of tipping points and regime shifts in coupled SESs [73] is gaining increasing interest [75], with many implications for the adaptive management of SESs [76]. Regime shifts can lead to social-ecological traps, where unsustainable feedbacks reinforce each others and push the SES into an undesirable state [9]. Some authors suggest that humanity may be locked in a technological innovation pathway that reinforces such unsustainable feedbacks [35]. Ecological time delays may also affect the human perception of environmental changes, thus worsening the amnesia and shortsightedness observed in conservation science, known as the shifting baseline syndrome [77]. This syndrome refers to a shift over time in the expectation of what a healthy biodiversity baseline is, and can lead to tolerate incremental loss of species through inappropriate management [78]. Time delays can also be related to perceived environmental uncertainty, which has been shown to endanger the enforcement of cooperation in SESs with common-pool resources [79].

Our results highlight the importance of accounting for the feedback loop between human demography, environmental degradation and behavioral changes when studying the long-term sustainability of coupled SESs. Especially, the temporal dynamics of coupled social-ecological processes matter, since ecological lag effects alter the human perception of environmental degradation and the rapidity of behavioral changes. Policies that enhance the adaptive capacity of social-ecological systems may thus benefit from taking social norms into account [8]. These insights also point to future research needs regarding the interplay of social, demographic and ecological long-term dynamics.

## Authors’ contributions

A.-S.L., C. de M. and M.L. jointly designed the study, developed the model and analyzed and interpreted the model results. A.-S.L. drafted the manuscript; M.L. and C. de M. revised it critically. All authors gave final approval for publication.

## Acknowledgements

We thank Matthieu Barbier, Kirsten Henderson and David Shanafelt for valuable discussions and helpful comments on earlier versions of the manuscript.

## 4 Electronic supplementary material

### 4.1 Functions and aggregate parameters

**Table S1:** Functions and aggregate parameters expression and definition. *i* = {*u, s*}; H: units of labor; t: units of time.

| Functions and aggregate parameters |  | Definition | Units |
| --- | --- | --- | --- |
| $\phi_i$ | $\kappa/(1 - \alpha_i\eta - \alpha_2(1 - \eta))$ | Population density on converted land | H |
| $S(H, q)$ | $\left(1 - \frac{H}{\phi(q)}\right)^z$ | Long-term species richness | — |
| $\gamma_2$ | $(1 - \eta) T_m (\alpha_2)^{\alpha_2} \left(\frac{1 - \alpha_2}{\kappa}\right)^{1 - \alpha_2}$ | Max. *per capita* industrial consumption | H<sup>-1</sup> |
| $\gamma_{1i}$ | $\eta T_m (\alpha_i)^{\alpha_i} \left(\frac{1 - \alpha_i}{\kappa}\right)^{1 - \alpha_i}$ | Max. *per capita* agricultural consumption | H<sup>-1</sup> |
| $y_{1i}$ | $\gamma_{1i} B^\Omega T/T_m$ | Mean *per capita* agricultural consumption | H<sup>-1</sup> |
| $y_2$ | $\gamma_2 T/T_m$ | *Per capita* industrial consumption | H<sup>-1</sup> |
| $U_i$ | $y_{1i}^\eta y_2^{1 - \eta}$ | *Per capita* consumption utility | H<sup>-1</sup> |
| $\Delta$ | $\epsilon - 4\Omega z y_1^{min} \left( \left( \frac{\gamma_{1u}}{y_1^{min}} \right)^{\frac{1}{\Omega z}} - 1 \right) e^{-b_2 \gamma_2} \mu$ | Sustainability criterion | t<sup>-1</sup> |

### 4.2 Parameters definition, units and defaults values

**Table S2:**
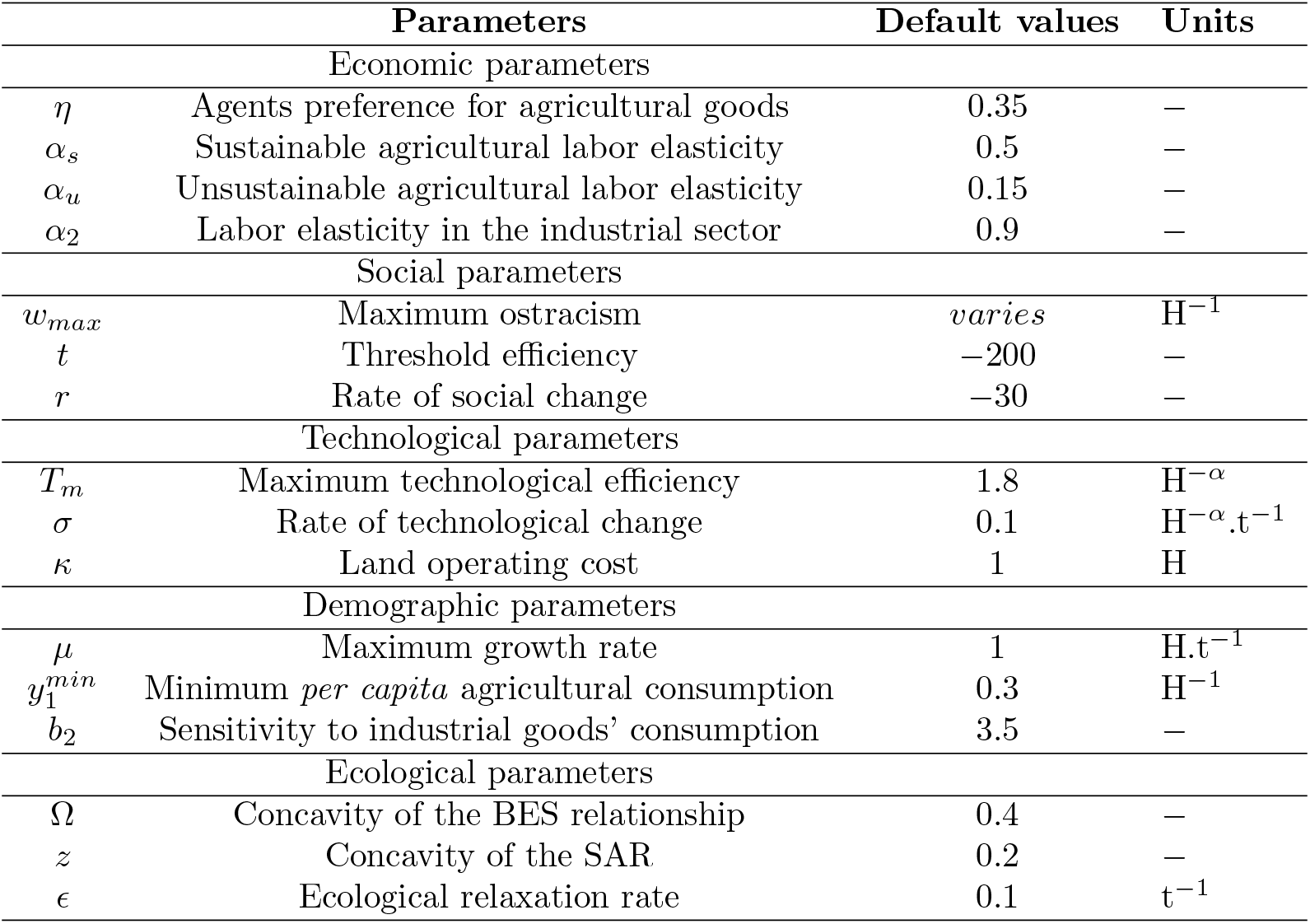
Definition, units and default values of the parameters. H: units of labor; t: units of time.

### 4.3 Dynamical system analysis

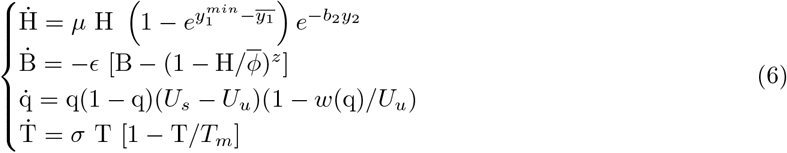

Parameters and functions are summarized in Tables S1 and S2, with 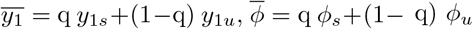, the consumption utility 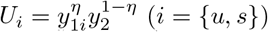, and the ostracism function *w*(*q*) = *w_max_e^te^rq^^*.

Solving system (6) for Ḣ = 0, Ḃ = 0, Ṫ = 0 and 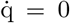 gives five equilibria: (1) a sustainable equilibrium, 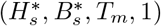, (2) an unsustainable equilibrium, 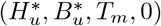, (3) a mixed equilibrium, 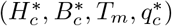, and (4) two unviable equilibria, (0, 1, *T_m_*, 0) and (0, 1, *T_m_*, 1).

We first evaluate the Jacobian matrix at the viable equilibria, (*H*\*, *B*\*, *q*\*, T_m_) where *q*\* = 1 or *q*\* = 0. After simplification, we obtain:

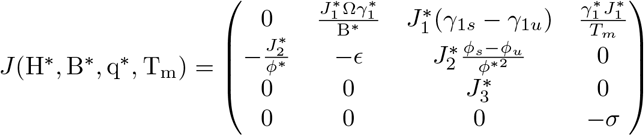

where 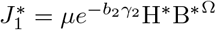, 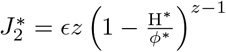, and 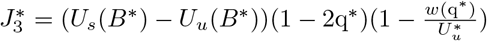.

The determinant *D* of this Jacobian matrix is the product of the four eigenvalues of the system. An equilibrium is locally stable if all its eigenvalues are negative, i.e. *D* > 0. In order to assess the local stability of the viable equilibria, lets first derive the determinant of *J*(H*, B*, q*, T_m_):

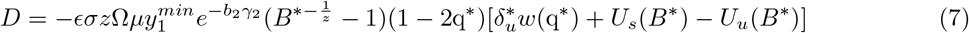

where 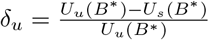.

We obtain the determinant *D_s_* of the Jacobian evaluated at the sustainable equilibrium by taking 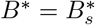 and *q*\* = 1, so that:

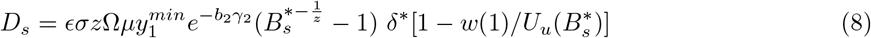

where 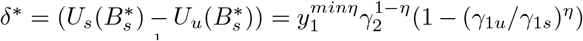. Since *γ*_1*u*_ > *γ*_1*s*_ and 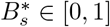, we deduce that *δ*\* < 0 and 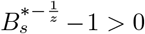, so that the sign of *D_s_* depends on the last term of eq. (8). The sustainable equilibrium 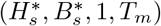 is thus locally stable (*D_s_* > 0) if

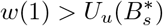

where 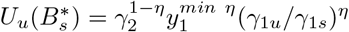.

The determinant *D_u_* of the Jacobian evaluated at the unsustainable equilibrium (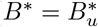 and *q*\* = 0 is:

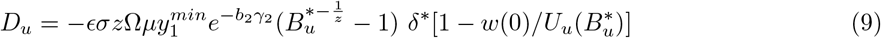

where 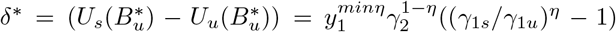. Since *γ*_1*u*_ > *γ*_1*s*_ and 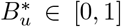, we deduce that *δ*\* < 0 and 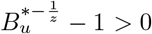, so that the sign of *D_u_* depends on the last term of eq. (8). The unsustainable equilibrium 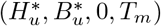 is thus locally stable (*D_u_* > 0) if

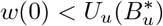

where 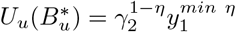.

Let us now evaluate the Jacobian matrix at the unviable equilibria, (0,1, q*, T_m_) where *q*\* = 1 or *q*\* = 0. After simplification, we obtain:

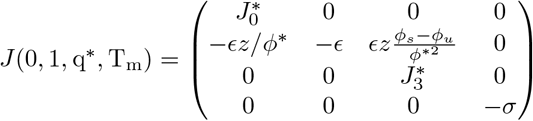

where 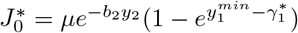.

The determinant *D*(0, 1, *q*\*, *T_m_*) writes:

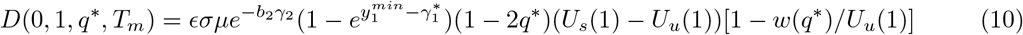

where 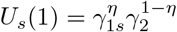 and 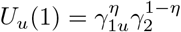, so that *U_s_*(1) < *U_u_*(1).

Therefore, the determinant *D*(0, 1, 0, *T_m_*) is:

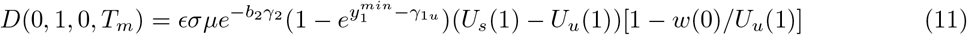

Thus, when the viable equilibria are feasible, i.e. when 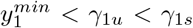, the unviable equilibrium (0, 1, 0, *T_m_*) is stable if *w*(0) > *U_u_*(1). However, since *U_u_*(1) > *U*\*, the unviable equilibrium (0, 1, 0, *T_m_*) is only stable when the corresponding viable equilibrium 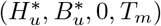 is unstable.

Similarly, the determinant D(0, 1, 1, *T_m_*) is:

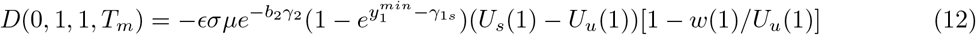

When the viable equilibria are feasible, the unviable equilibrium (0, 1, 1, *T_m_*) is stable if *w*(1) < *U_u_*(1). In this case, both viable 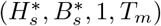 and unviable (0, 1, 1, *T_m_*) equilibria can be stable at the same time, if 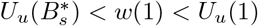.

